# The Aryl Hydrocarbon Receptor Promotes Differentiation During Mouse Preimplantational Embryo Development

**DOI:** 10.1101/2020.04.07.026047

**Authors:** Ana Nacarino-Palma, Jaime M. Merino, Pedro M. Fernández-Salguero

**Author notes:** Corresponding authors: Pedro M. Fernández-Salguero, Jaime M. Merino. **Abbreviations:** AhR, Aryl Hydrocarbon Receptor; ICM, internal cell mass; TE, trophectoderm; TMRM, Tetramethylrhodamine; YAP, Yes-activated protein.

## Abstract

Mammalian embryogenesis is a complex process controlled by transcription factors that dynamically regulate the balance between pluripotency and differentiation. Transcription factor AhR is known to regulate *Oct4/Pou5f1* and *Nanog*, both essential genes in pluripotency, stemness and early embryo development. Yet, the molecular mechanisms controlling *Oct4/Pou5f1* and *Nanog* during embryo development remain largely unidentified. Here, we show that AhR is required for proper embryo differentiation by regulating pluripotency factors and by maintaining adequate metabolic activity. AhR lacking embryos (*AhR*-/-) showed a more pluripotent phenotype characterized by a delayed expression of differentiation markers of the first and second cell divisions. Accordingly, central pluripotency factors OCT4/POU5F1, NANOG, and SOX2 were overexpressed in *AhR*-/- embryos at initial developmental stages. An altered intracellular localization of these factors was observed in absence of AhR and, importantly, OCT4 had an opposite expression pattern with respect to AhR from the 2-cell stage to blastocyst, suggesting a negative regulatory mechanism of OCT4/POU5F by AhR. Hippo signalling, rather than being repressed, was upregulated in very early *AhR*-/- embryos, possibly contributing to their undifferentiation at later stages. Consistently, AhR-null blastocysts overexpressed the early marker of inner cell mass (ICM) differentiation Sox17 whereas downregulated extraembryonic differentiation-driving genes *Cdx2* and *Gata3*. Moreover, the persistent pluripotent phenotype of *AhR*-/- embryos was supported by an enhanced glycolytic metabolism and a reduction in mitochondrial activity. We propose that AhR is a regulator of pluripotency and differentiation in early mouse embryogenesis and that its deficiency may underline the reduced viability and increased resorptions of AhR-null mice.

## INTRODUCTION

The aryl hydrocarbon receptor (AhR) is a transcription factor with important toxicological and physiological implications and roles in pluripotency and stemness recently identified (Ko and Puga, 2017; Mulero-Navarro and Fernandez-Salguero, 2016; Roman et al., 2018). Several studies support this receptor as an important regulator of the balance between pluripotency and differentiation under physiological conditions and in tumor cells. Indeed, AhR activation by the carcinogen TCDD during mouse pregnancy blocked the ability of hematopoietic stem cells (HSC) for long-term self-renewal (Laiosa et al., 2015). Similarly, sustained AhR activation during early differentiation of mouse embryonic stem cells impaired signalling critical for the ontogeny of cardiac mesoderm and cardiomyocyte functions (Wang et al., 2016). Previous work from our laboratory using human NTERA-2 cells revealed that AhR supports cell differentiation through the transcriptional repression of retrotransposable *Alu* elements located in the promoter region of pluripotency genes *Oct4* and *Nanog* (Morales-Hernandez et al., 2016). On the contrary, receptor deficiency in mice produces a more undifferentiated phenotype improving the regenerative potential of the lung (Morales-Hernandez et al., 2017) and the liver (Moreno-Marin et al., 2017) upon acute damage.

A distinguishing feature of preimplantation development is the gradual loss of totipotence of the embryonic stem cells (ESCs). Throughout embryonic development from zygote to blastocyst, ESCs will restrict their fate through cellular differentiation after successive rounds of cell division. Three different cell lineages exist in the mature blastocyst; namely, trophectoderm, epiblast and primitive endoderm (Chazaud and Yamanaka, 2016). In the first cell fate decision, asymmetric divisions in the initial embryo generate outside and inside cells that differ in their cellular properties, location within the embryo and cell outcome (Fleming, 1987; Johnson and Ziomek, 1981; Morris et al., 2010). Outside cells will differentiate into the trophectoderm (TE), which is the precursor lineage of the placenta. Inside cells constitute the pluripotent inner cell mass (ICM) that will differentiate in the second cell fate decision to form the primitive endoderm (PE) giving rise to the yolk sac, and the pluripotent embryonic epiblast (EPI) that is the precursor of all embryonic tissues. Numerous signalling networks are responsible for coordinating the myriad events needed to control the balance between differentiation and pluripotency in embryogenesis. Transcription factors OCT4/POU5F1 (herein OCT4), SOX2 and NANOG constitute the “central pluripotency network” (Boyer et al., 2005; Boyer et al., 2006). These pluripotency factors are initially expressed in all cells of the morulae, with their expression becoming gradually restricted to the ICM after first cell fate decision (Bedzhov et al., 2014). Establishment of TE fate program in outside cells is regulated by the Hippo pathway, which acts as a sensor of cell polarity.

Outside cells have asymmetric cell-cell contacts that lead to the accumulation of apical polarity proteins that inhibit activity of the tight junction proteins AMOT and the Hippo pathway kinases LATS1/2 (Leung and Zernicka-Goetz, 2013; Paramasivam et al., 2011). As a result, hypophosphorylated Yes-activated protein (YAP) is translocated to the nucleus and the TE cell fate program is activated with an increase in CDX2 expression through TEAD4. In inside cells, symmetric cell-cell contacts prevent the establishment of an apical domain. AMOT proteins are then activated and distributed by all membrane in adherent junctions in a NF2/α-Catenin/β-catenin/E-cadherin complex. In addition, LAST1/2 become activated and the resulting phosphorylated YAP excluded from the nucleus; OCT4 is then expressed and the pluripotency program initiated to determine the ICM fate (Manzanares and Rodriguez, 2013).

Knowing how cell fate is specified in the preimplantation embryo may help to understand the mechanisms that regulate pluripotency and differentiation of stem cells of embryonic origin as well as those arising from tumors.

Interestingly, early and previous reports have shown that AhR-null mice have a reduced fertility producing fewer numbers of pups born alive as compare to AhR-expressing littermates. In fact, such phenotype seems to be at least partially due to an increase in embryo resorption and to an impaired ability to complete the preimplantation program to the blastocyst stage (Abbott et al., 1999; Peters and Wiley, 1995). Here, we have studied how AhR affects the early stages of preimplantation during mouse development in an attempt to further understand receptor functions in pluripotency and differentiation. We have found that AhR has pro-differentiation functions in the early mouse embryo needed to specify the different cell fates from the one-cell to the blastocyst stage. Our results suggest that AhR has relevant roles in embryonic stem cell differentiation through the control of genes responsible for maintaining a pluripotent status. AhR deficiency may thus negatively affect embryo progression during preimplantation eventually compromising viability.

## RESULTS

## AhR expression and localization is modulated throughout embryonic development

To analyze the role of AhR in early embryo differentiation, we first analyzed AhR expression levels along different embryonic stages. Confocal immunofluorescence analysis showed that AhR was significantly and steadily expressed as differentiation progressed from 2-cell zygote to blastocyst **(Fig. 1A,B)**. Regarding AhR localization within the embryo, the immunofluorescence analysis revealed a generalized expression in all cells up to the morulae stage with some cells having nuclear AhR. However, as differentiation progressed to the early and late blastocyst, AhR was only detected in the external blastomeres being almost absent in those cells forming the ICM **(Fig. 1A)**. To further support this finding, we separated inner and outer (TE) blastomeres from blastocysts using magnetic-activated cell sorting and analyzed AhR expression in both fractions. The results confirmed that AhR mRNA levels were significantly higher in TE blastomeres than in inner cell mass blastomeres **(Fig. 1C)**. Consistently, AhR mRNA expression significantly increased during differentiation from zygote to blastocyst at the transcriptional level **(Fig. 1D)**. These results indicated that the expression of the aryl hydrocarbon receptor is modulated throughout early embryonic development and that its embryonic localization changes with differentiation.

**Figure 1.**
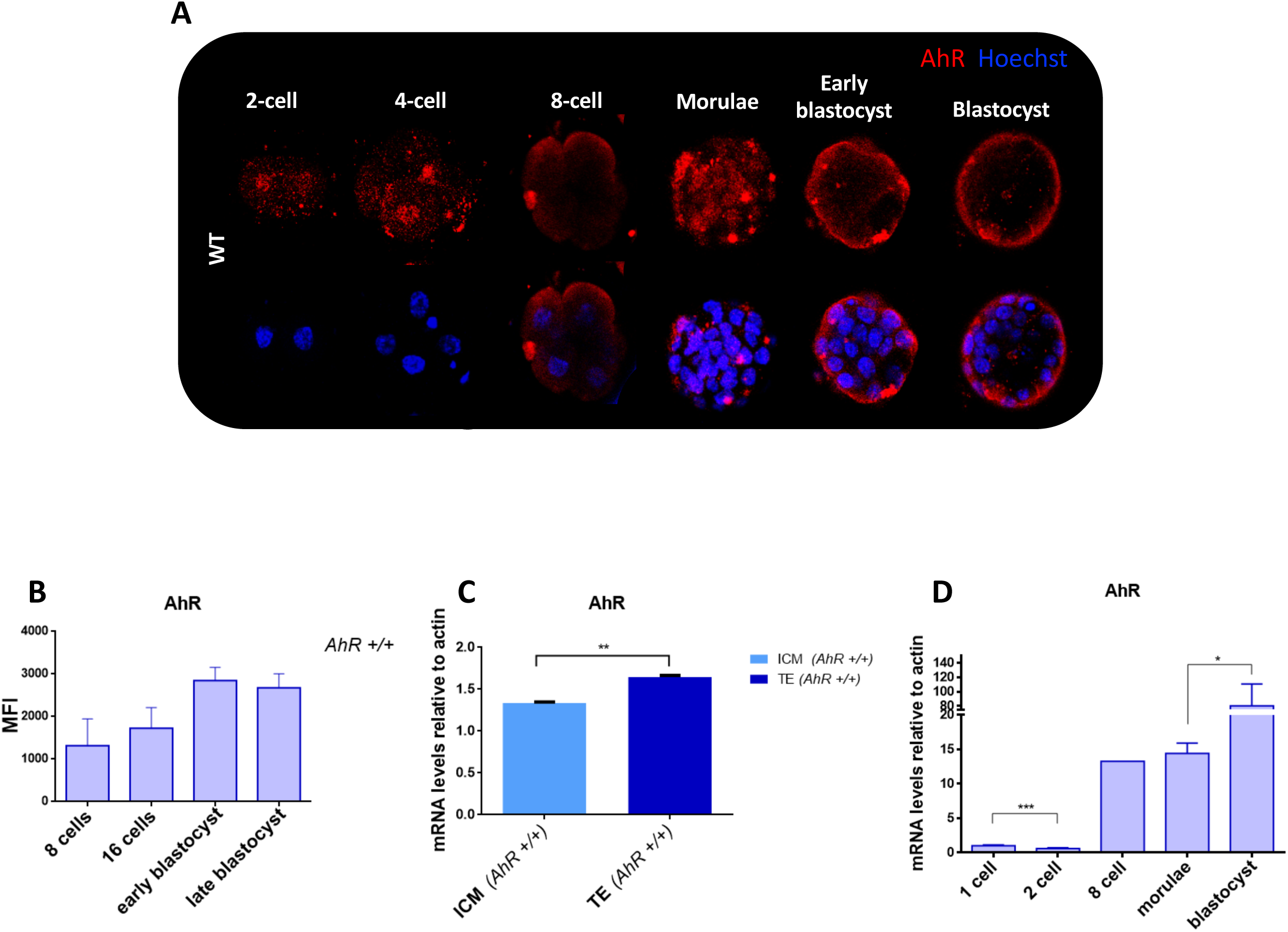
AhR expression increases during embryo differentiation. **(A)** Inmmunofluoresence analysis of AhR at the indicated embryonic stages. Whole *AhR*+/+ embryos were stained using a specific AhR antibody. Hoechst staining was used to label the cell nucleus. Confocal microscopy was used for detection. **(B)** Immunofluorescence was quantified by calculating the mean fluorescence intensity (MFI) for each develpmental stage. **(C)** *AhR* mRNA expression was quantified by RT-qPCR using RNA purified from TE or ICM fractions previously separated by MACS. **(D)** *AhR* mRNA expression was quantified by RT-qPCR in *AhR+/+* embryos at the indicated developmental stages using total RNA and the specific primers indicated in Supplementary Table S1. RT-qPCR was normalized by the expression of *β*-*Actin* and represented as 2^−ΔΔCt^. \**p*< 0.05; \*\**p*< 0.01. Data are shown as mean ± SD.

### AhR deficiency induces upregulation of pluripotency genes during early embryo development

In order to assess whether the aryl hydrocarbon receptor participates in the maintenance of pluripotency during the early stages of embryogenesis, we next analyzed the levels of pluripotency factors throughout preimplantation in wild type and AhR-null embryos. *AhR*-/- embryos showed significantly higher *Nanog* and *Oct4* mRNA levels as compared to *AhR+/+* embryos from 1-cell zygote until the morulae stage **(Fig. 2A,B)**. In blastocysts, *Oct4* expression kept rising in AhR-null embryos while *Nanog* mRNA levels became balanced among both genotypes **(Fig. 2A,B)**. Regarding *Sox*2, embryos lacking AhR also had higher expression of this pluripotency factor at the beginning of embryogenesis, 1-cell and 2-cell stages, to decrease to similar levels in both genotypes from 8-cell to blastocyst **(Fig. 2C)**. Thus, AhR plays a role in controlling the expression of genes known to regulate pluripotency and the differentiation required for embryo development.

**Figure 2.**
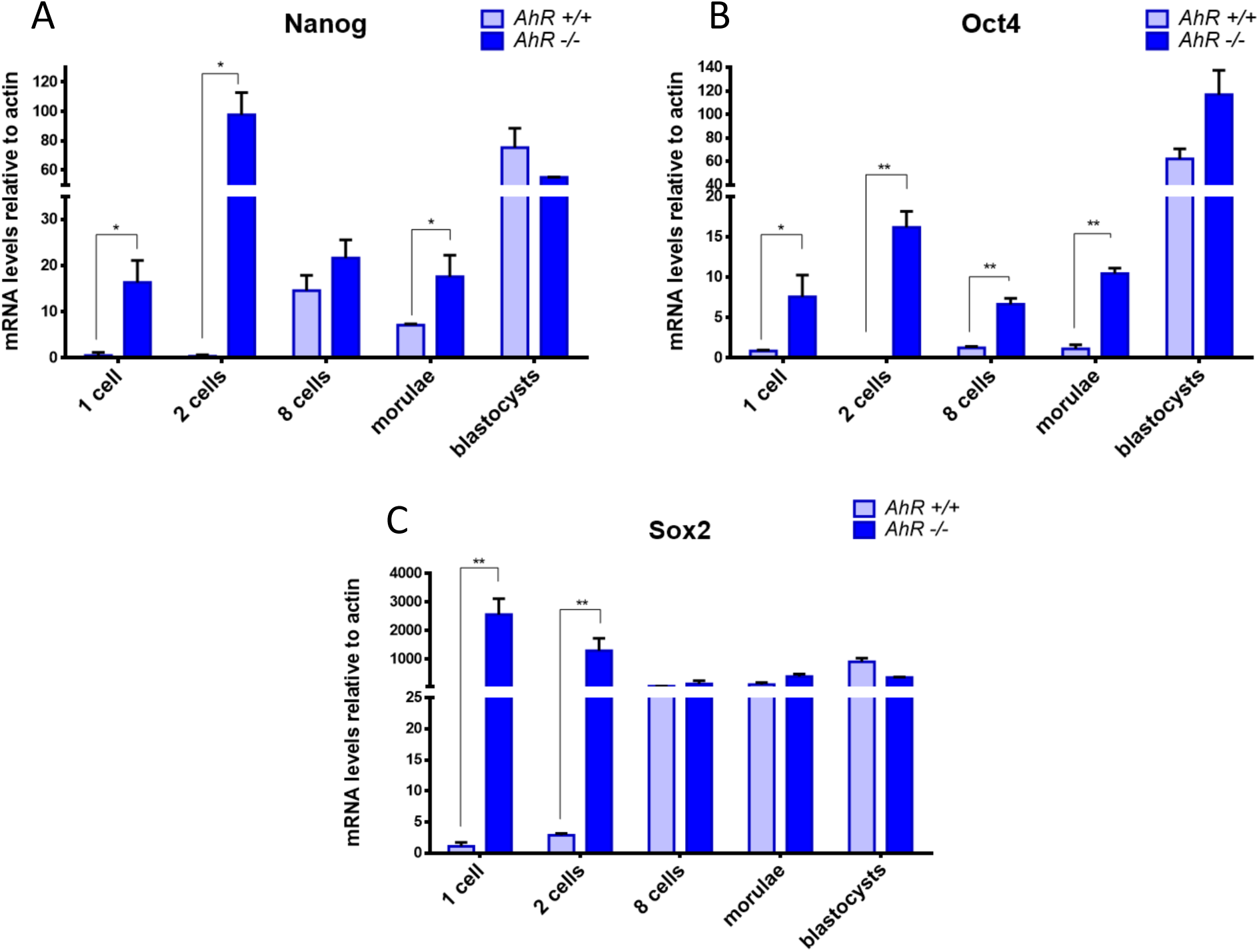
Pluripotency factors are upregulated in AhR-null embryos. **(A-C)** *AhR+/+* and *AhR*-/- embryos were obtained at the indicated embryonic stages and used to quantify the mRNA expression of *Nanog* **(A)**, *Oct4* **(B)**, *and Sox2* **(C)** by RT-qPCR. Expression levels were normalized by *β*-*Actin* and represented as 2^−ΔΔCt^. \**p*< 0.05; \*\**p*< 0.01. Data are shown as mean ± SD.

### AhR modulates the localization of pluripotency factors during embryogenesis

To investigate how AhR affects protein levels and localization of pluripotency factors, we did immunofluorescence analysis for OCT4 and NANOG in wild type and AhR-null embryos during blastocyst development. The results obtained showed changes in OCT4 and NANOG localization upon the presence of the aryl hydrocarbon receptor. In wild type embryos, OCT4 had a preferred cytoplasmic localization up to the early blastocyst stage, to then move to the nucleus in cells at the ICM **(Fig. 3A)**. Embryos lacking AhR showed nuclear localization of OCT4 throughout most stages of development and up to late blastocyst **(Fig 3A)**. We then decided to analyze OCT4 expression at the mRNA level in isolated blastomeres from the ICM and TE of both genotypes. We found that in *AhR*-/- embryos, there were no significant differences in OCT4 mRNA expression between ICM and TE blastomeres, as it was observed in *AhR+/+* embryos **(Fig 3B)**. These results indicate that AhR may be needed to regulate the location of OCT4 within the embryo, which could impact differentiation and cell fate. The fact that OCT4 has a location pattern opposite to that of AhR **(Figs. 1 and 3C)**, suggests that AhR may exert a negative regulation on OCT4 to drive embryo differentiation.

**Figure 3.**
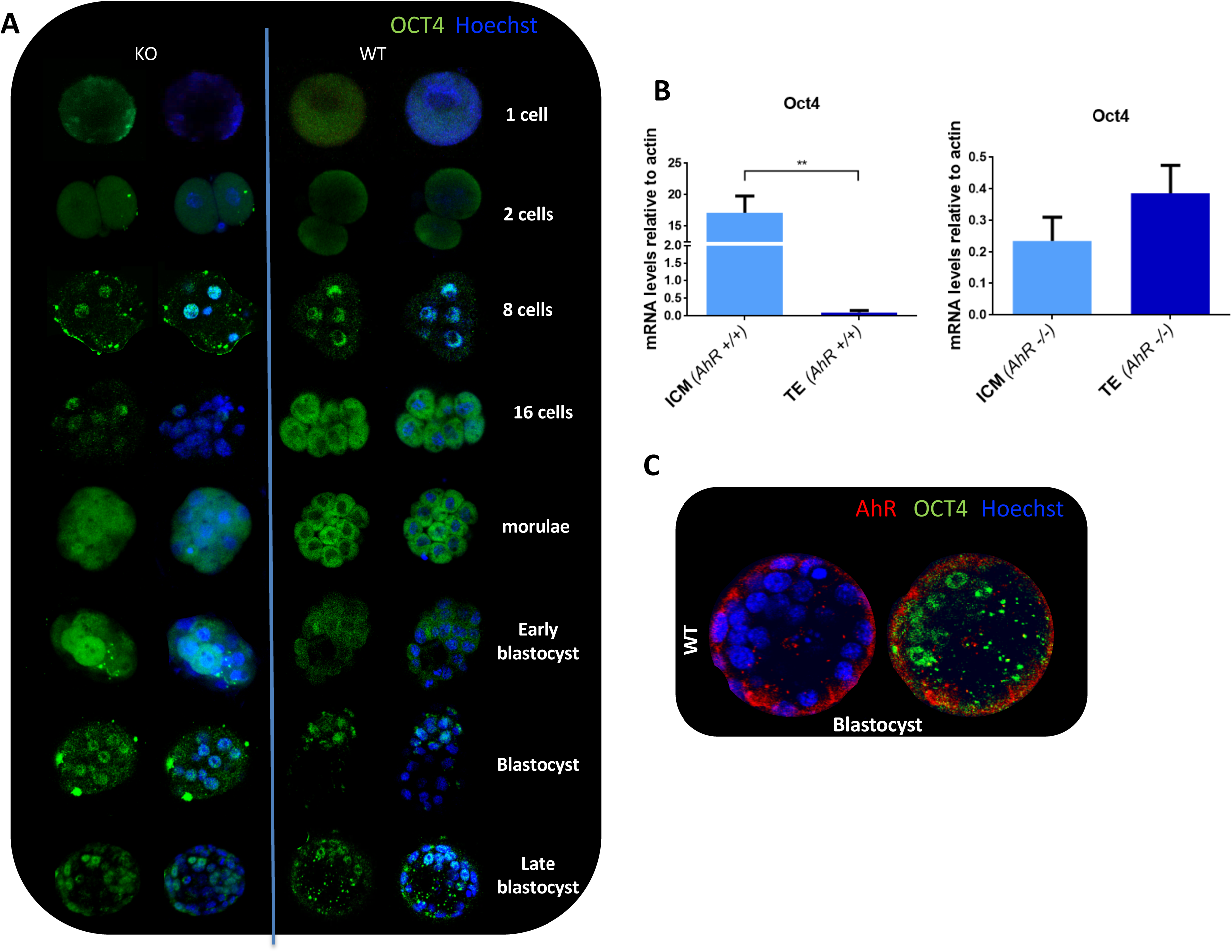
AhR depletion alters OCT4 cellular distribution through embryogenesis. **(A)** Inmmunofluoresence analysis of OCT4 at the indicated embryonic stages. Whole embryos were stained using a specific antibody. Hoechst was used to stain cell nuclei. **(B)** *Oct4* mRNA expression was quantified by RT-qPCR using mRNA purified from TE and ICM fractions separated by MACS using the specific primers indicated in Supplementary Table S1. (**C)** Inmmunofluoresence analysis of OCT4 (green) and AhR (red) in embryos at the blastocyst stage. mRNA expression was normalized by *β*-*Actin* and represented as 2^−ΔΔCt.^. \*\**p*< 0.01. Data are shown as mean ± SD. Confocal microscopy was used for detection.

NANOG localization in *AhR+/+* embryos was also modified in the absence of AhR. The dotted and regular pattern that this protein had from zygote to 4-cell in AhR wild type embryos, remained in AhR-null embryos until the 16-cell stage **(Fig 4)**. While in AhR wild type embryos a polarized and nuclear distribution of NANOG between ICM and TE was observed from moruale on, a delocalization of this pluripotency factor was evident in AhR lacking embryos **(Fig 4)**. Quantification of the immunofluorescence signals revealed that global NANOG expression was significantly higher in *AhR*-/- than in *AhR+/+* embryos **(Fig. 4B)**. These data suggest that, in addition to OCT4, AhR could also regulate NANOG expression during embryo development.

**Figure 4.**
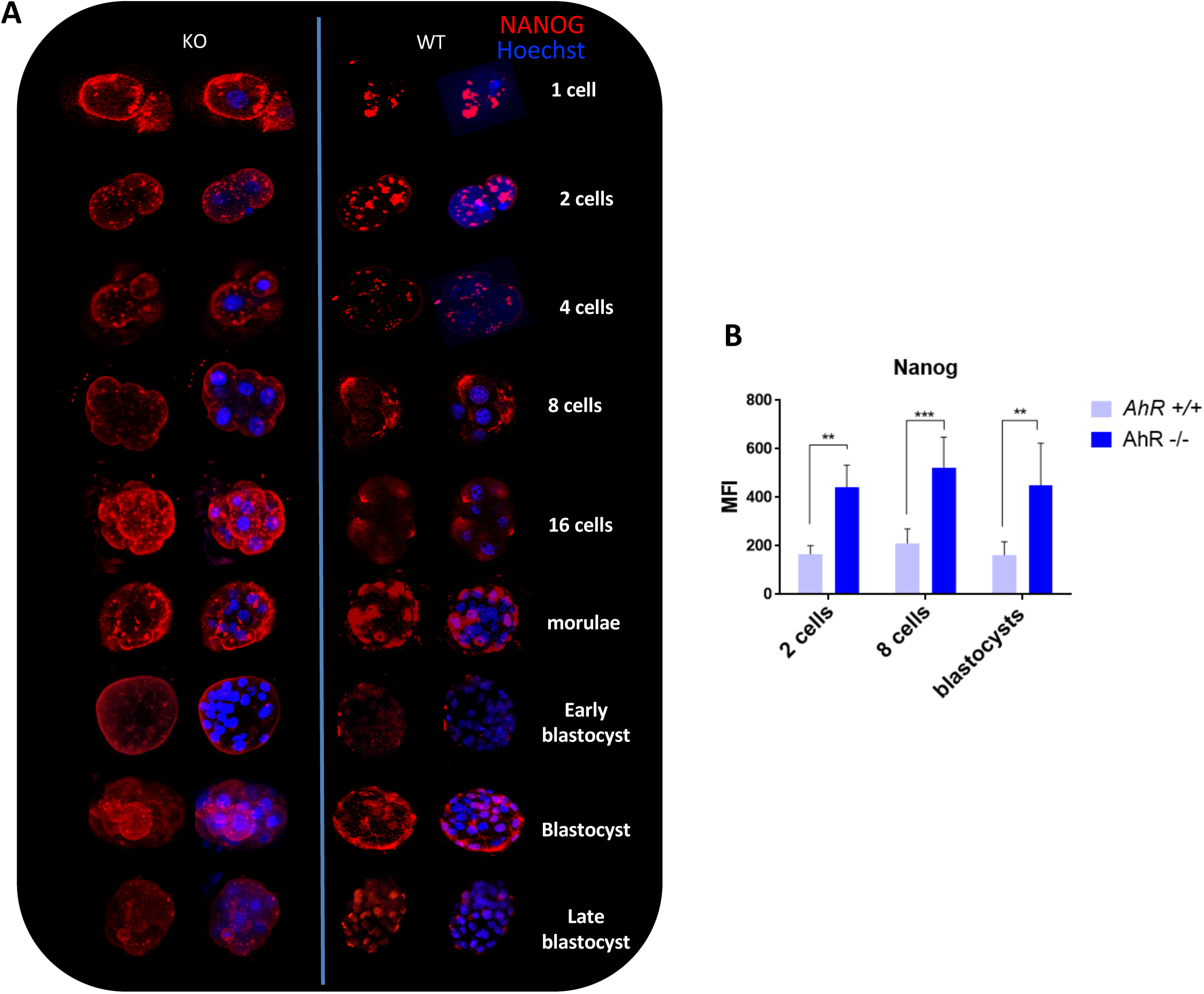
NANOG distribution in the embryo is altered in absence of AhR. **(A)** Inmmunofluoresence analysis of NANOG at the indicated embryonic stages. Whole embryos were stained using a specific antibody. Hoechst was used for staining cell nuclei. **(B)** Immunofluorescence was quantified by calculating the mean fluorescence intensity (MFI). \*\**p*< 0.01; \*\*\**p*< 0.001. Data are shown as mean ± SD.

### AhR-null embryos show Hippo signalling upregulation

As indicated above, the Hippo pathway is implicated in cell polarity and cell fate. Next, we explored if the effects of AhR on embryonic differentiation could be mediated through the Hippo pathway. To investigate such possibility, we first analyzed the nuclear localization of the Hippo effector YAP. YAP was excluded from the cell nucleus in AhR-null embryos during most of embryo development, whereas it was located in the nucleus of the external blastomeres in wild type embryos from the morulae stage **(Fig 5A)**. Immunofluorescence analysis indicated that pYAP was predominantly excluded from the cell nucleus in a fraction of blastomeres in AhR-null blastocysts **(Fig. 5B)**. Quantification of the mean fluorescence intensity (MFI) revealed that pYAP levels (e.g. cytosolic) were significantly higher in *AhR*-/- embryos **(Fig. 6A)** and, consequently, that the amounts of nuclear YAP (unphosphorylated) were reduced in absence of AhR **(Fig. 6B)**.

**Figure 5.**
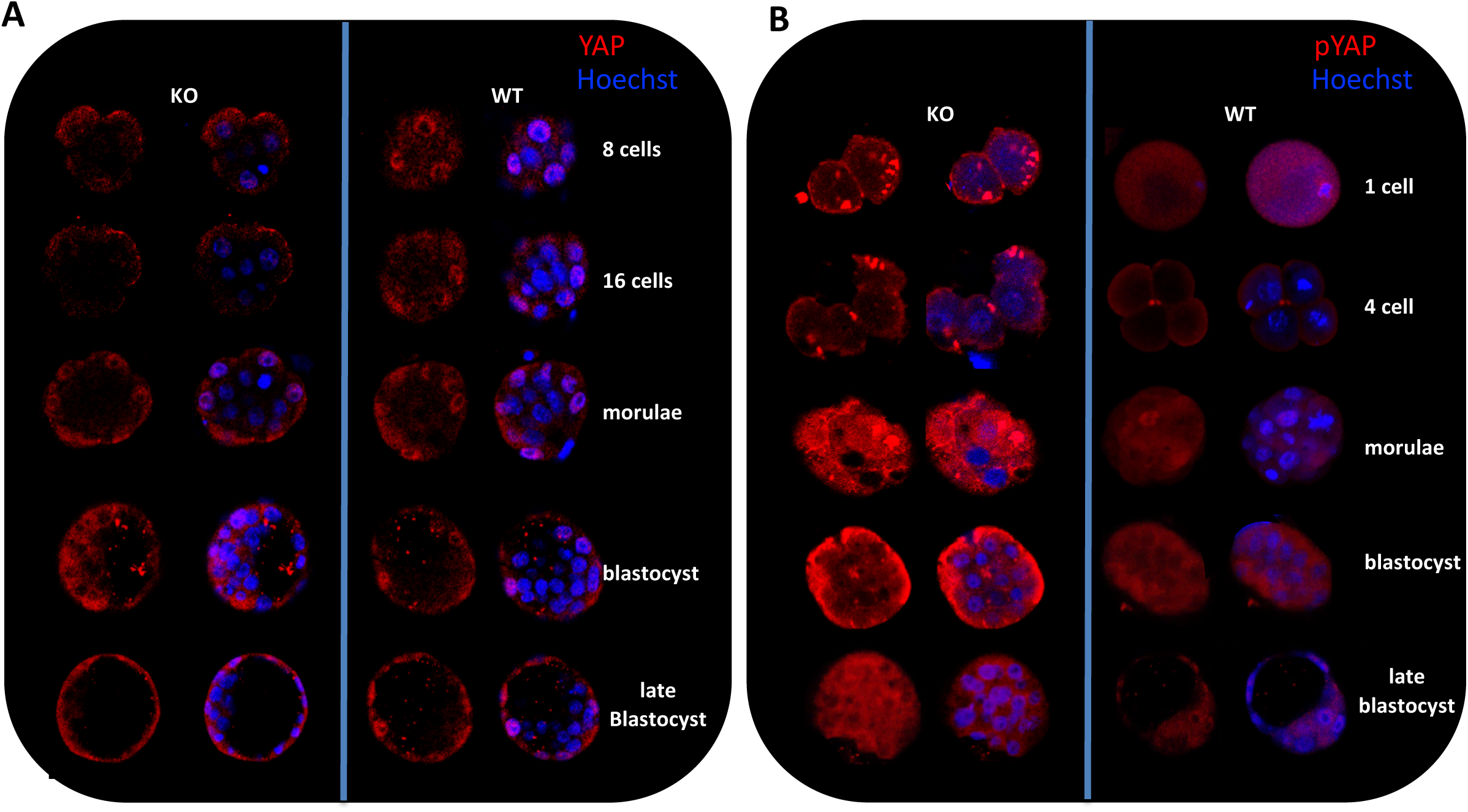
AhR deficiency alters localization of the Hippo effector YAP. Inmmunofluoresence analysis of YAP and pYAP at the indicated developmental stages in *AhR+/+* and *AhR*-/- embryos **(A,B)**. Whole embryos were stained using specific antibodies for YAP **(A)** or phosphor-YAP **(B)**. Hoechst was used for staining of cell nucleus. Confocal microscopy was used for detection.

**Figure 6.**
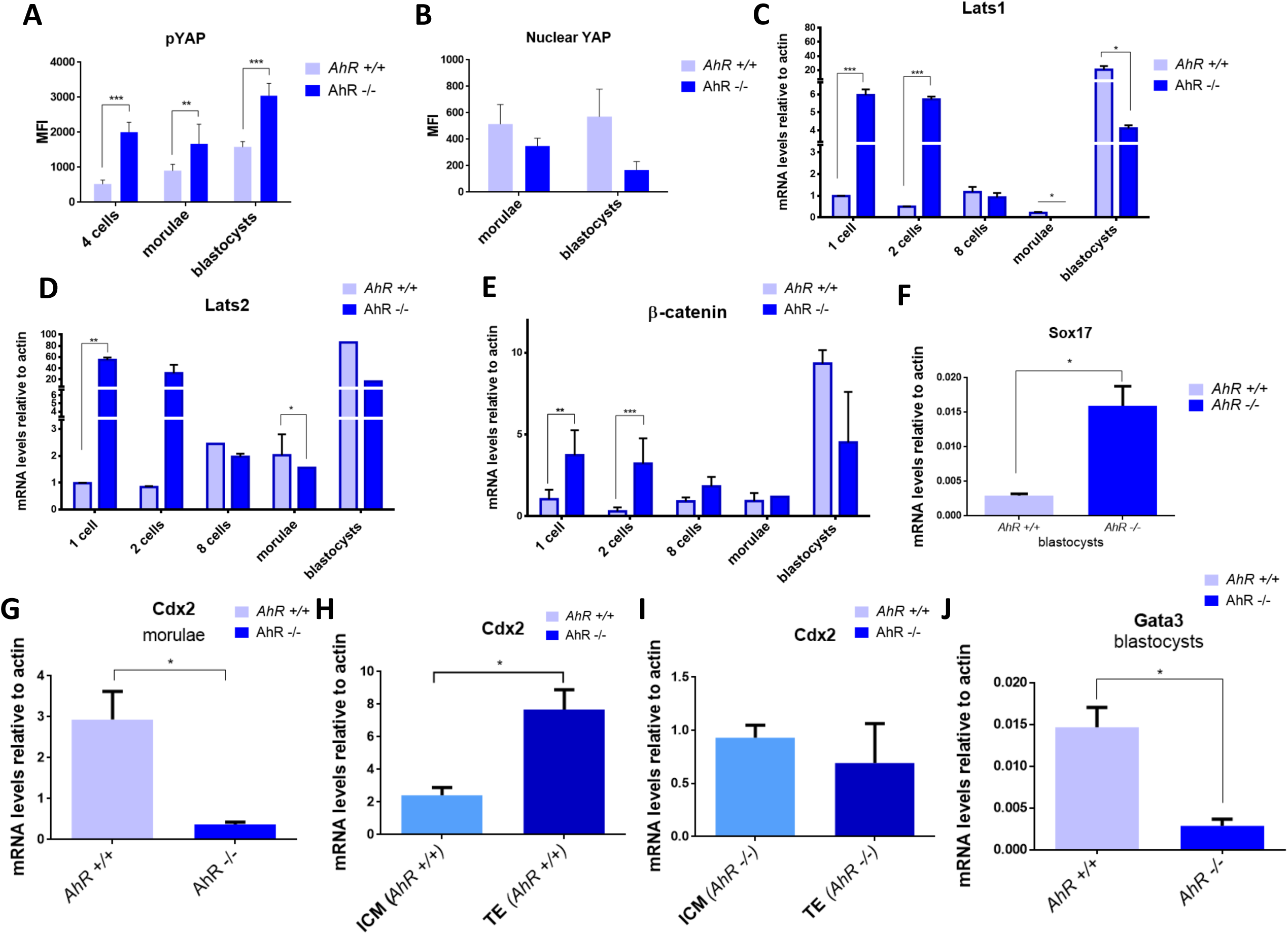
Hippo effectors and molecular intermediates of pluripotency are altered in absence of AhR. **(A)** Levels of pYAP were quantified from immunofluorescences and represented as the mean fluorescence intensity (MFI). **(B)** Nuclear YAP levels in morulae and blastocysts from *AhR+/+* and *AhR*-/- mice was quantified by immunofluorescence and represented as the mean fluorescence intensity (MFI). *AhR+/+* and *AhR*-/-embryos at the indicated developmental stages were used to purify embryo mRNA **(C-G, J)** or mRNA from TE and ICM fractions **(H,I)** that were separated by MACS as indicated in Material and Methods. The expression of *Lats1* **(C)**, *Lats2* **(D)**, *β-Catenin* **(E)**, *Sox17* **(F)**, *Cdx2* **(G-I)** and *Gata3* **(J)** was quantified by RT-qPCR. Expression levels were normalized by *β-Actin* and represented as 2^−ΔΔCt^. \**p*< 0.05; \*\**p*< 0.01; \*\*\**p*< 0.001. Data are shown as mean ± SD.

To further analyze the implication of the Hippo pathway, we measured the expression of the kinases responsible for YAP phosphorylation *Lats1* and *Lats2*. The results showed that their expression was significantly higher at the beginning of development (1-cell and 2-cell) in embryos lacking the aryl hydrocarbon receptor than in wild type ones **(Fig 6C,D)**. Interestingly, the levels of both kinases transiently decreased from 8-cell to morulae to increase again at the blastocyst stage **(Fig 6C,D)**. In addition, β-catenin, a component of the complex located at the adherent junctions where AMOT is retained, was overexpressed at the initial stages of development in *AhR*-/- embryos **(Fig. 6E)**. The early marker for ICM pluripotency and undifferentiation *Sox17*, was also overexpressed in *AhR*-/- blastocysts **(Fig. 6F)**, in agreement with previous studies indicating that Sox17 is expressed at ICM as a endoderm primitive marker (Artus et al., 2011; Frum and Ralston, 2015). Moreover, the transcriptional YAP target *Cdx2* was repressed in AhR-null morulae with respect to wild type morulae **(Fig. 6G)**. *Cdx2* expression was significantly higher in TE than in ICM from *AhR+/+* blastocysts **(Fig. 6H)** whereas no significant differences were found in *Cdx2* distribution between the ICM and TE in *AhR*-/- blastocysts **(Fig. 6I)**, further supporting an increased activation of the Hippo pathway, concomitant with a reduced transcriptional activity of the OCT4 repressor YAP, in absence of AhR. *Gata3*, a trophoectoderm marker in blastocysts, had lower levels in *AhR*-/- embryos at the blastocyst phase **(Fig. 6J)** supporting that lack of AhR promotes a more undifferentiated phenotype in preimplantation mouse embryos. Altogether, these data suggest that absence of AhR affects differentiation of the TE and ICM as the two first cell lineages established in the embryo.

### Embryos lacking AhR show a higher glycolytic metabolic activity and a lower rate of oxidative metabolism

The more undifferentiated status of *AhR*-/- embryos could be associated to a more immature physiological phenotype. We next decided to study glycolytic and oxidative metabolism rates since it is well established that these two parameters are strongly linked to the pluripotency state of embryonic stem cells. First, we analyzed the mitochondrial membrane potential of embryos of both genotypes at different stages using tetramethyl rhodamine (TMRM) staining. **E**mbryos lacking AhR maintained a lower mitochondrial activity until the 32-cell stage, while wild type embryos had a significantly higher mitochondrial activity during the same period **(Fig. 7A,B)**. To further asses this result, we collected pools of *AhR*-/- and *AhR+/+* embryos and analyzed their mitochondrial membrane potential using the JC-10 probe. The results confirmed that the mitochondrial membrane potential was higher in wild-type than in AhR-null embryos **(Fig. 7C)**. Moreover, the mitochondrial volume measured by mitotracker green staining was significantly lower in *AhR*-/- than in *AhR+/+* embryos **(Fig. 7D)**, as well as the mRNA levels of the marker for mitochondrial activity mitochondrial carrier homolog-1 (Mtch1) **(Fig. 7E)**. These results indicate that lack of AhR may contribute to a lower rate of oxidative metabolism in the mouse embryo.

**Figure 7.**
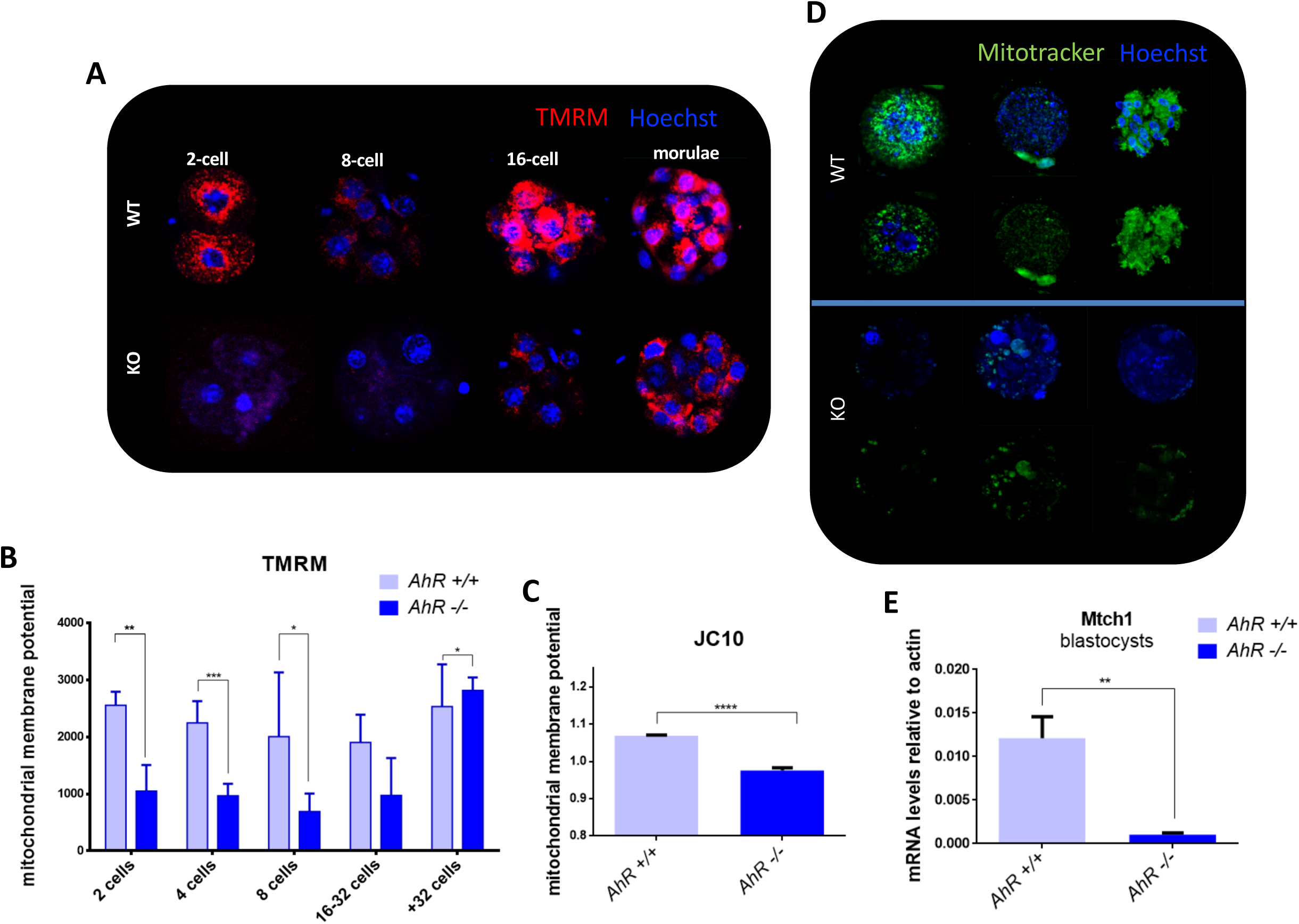
AhR lacking embryos have lower mitochondrial activity. Mitochondrial membrane potential was measured by TMRM staining at the indicated stages and then analyzed by confocal microscopy. Three embryos per genotype were analyzed (**A,B**). Mitochondrial membrane potential was determined by JC10 staining in pools of 15-20 embryos **(C)**. Mitochondrial volume was analyzed by mitotracker green staining and visualized by confocal microscopy **(D)**. Mitochondrial carrier homolog-1 (*Mtch1*) mRNA expression was measured in blastocysts fro each genotype. quantified by RT-qPCR. Expression levels were normalized by *β-Actin* and represented as 2^−ΔΔCt^. \**p*< 0.05; \*\**p*< 0.01; \*\*\**p*< 0.001. Data are shown as mean ± SD.

Next, we investigated if glycolytic metabolism, the preferred energy source for pluripotent and cancerous cells, would be influenced by AhR activity throughout embryonic differentiation. We observed that the expression of the hexokinase enzyme (*HK*) and of glucose transporters *Scl2a1* and *Scl2a3* were increased in *AhR*-/- as compared to *AhR+/+* blastocysts **(Fig. 8A-C)**. We then decided to measure hexokinase activity using an enzymatic assay in embryos at developmental stages between morulae and blastocyst. The results obtained revealed that absence of AhR generated a significant increase in hexokinase activity **(Fig. 8D**). The less differentiated status of AhR deficient embryos with respect to wild type ones correlates with their preferential glycolytic metabolism. Thus, lack of AhR alters mitochondrial functions that are consistent with a more pluripotent phenotype. Stem cells specifically use the amino acid threonine to maintain their pluripotent status and such cellular condition is dependent on the activity of the threonine dehydrogenase (TDH) (Wang et al., 2009). We have found that TDH expression was increased in absence of AhR along embryonic development from 1-cell zygote to blastocyst, supporting that AhR-null preimplantation embryos have al altered metabolic profile.

**Figure 8.**
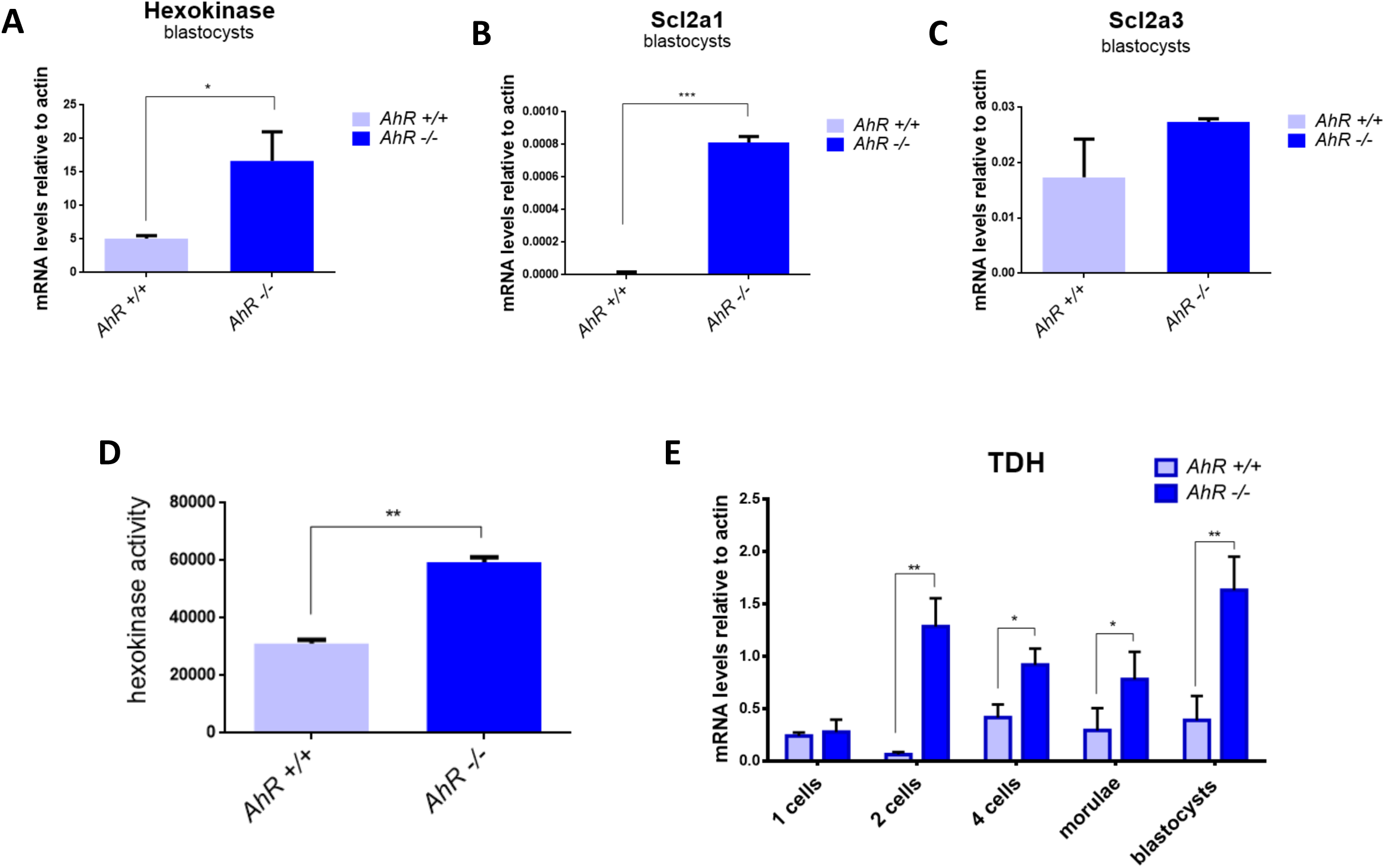
AhR depletion favors a glycolytic metabolism. mRNA was purified from *AhR+/+* and *AhR*-/- blastocysts and the expression of Hexokinase-*HK* **(A)**, *Scl2a1* **(B)** and *Scl2a3* **(C)** was quantified by RT-qPCR. Expression levels were normalized by *β-Actin* and represented as 2^−ΔΔCt^. **(D)** Hexokinase activity was measured using pools of 25-30 embryos at the morulae to blastocyst stages by an enzimatic assay **(E)**. Threonine dehydrogenase expression was analyzed at the indicated developmental stages in *AhR+/+* and *AhR*-/- embryos. \**p*< 0.05; \*\**p*< 0.01; \*\*\**p*< 0.001. Data are shown as mean ± SD.

## DISCUSSION

AhR promotes cell differentiation through the inhibition of pluripotency genes. As a result, AhR deficiency originates an undifferentiated phenotype not only in cell lines but also in tissue regeneration in mice (Morales-Hernandez, Gonzalez-Rico et al. 2016, Morales-Hernandez, Nacarino-Palma et al. 2017, Moreno-Marin, Barrasa et al. 2017). However, our knowledge about the role of AhR in embryo differentiation is still very limited, in particular with respect to the molecular intermediates that, been dependent on AhR, may be involved. This encouraged us to investigate the role of AhR in ESCs differentiation *in vivo* during mouse preimplantation embryonic development. In this phase of embryogenesis, totipotent blastomeres generate the first three cell lineages of the embryo: trophectoderm, epiblast and primitive endoderm. Mouse embryogenesis has been widely studied to understand developmental processes in mammals, but it also constitutes an excellent model to study the plasticity of stem cells. Understanding how molecular intermediates govern the balance between pluripotency and differentiation in blastocyst development allows us to understand stem cell behavior in other physiological and pathological conditions. An important finding of this study is that AhR affects the differentiation processes of embryo development by interacting with different signaling networks.

As cleavage of the early zygote takes place, central pluripotency factors increase their expression to produce totipotent cells that will proliferate and differentiate to generate a complete organism. We have first found that zygotes from *AhR*-/- mice have basal overexpression of well-known pluripotency factors *Oct4, Nanog* and *Sox2*, in agreement with our previous studies showing that AhR-null mice have an increased ability to regenerate lung (Morales-Hernandez et al., 2017) and liver (Moreno-Marin et al., 2017) and a higher potential to sustain undifferentiation of human embryionic carcinoma cells (Morales-Hernandez et al., 2016). The apparent global role of AhR in controlling differentiation was also reported by its ability to regulate ovarian follicular development through piRNA-associated proteins, piRNAs and retrotransposons (Rico-Leo et al., 2016).

Pluripotency genes need to reach a certain expression level to activate the networks that control pluripotency. Interestingly, *Oct4, Nanog and Sox2* reached their highest expression levels in a transient manner in 1-cell and 2-cell *AhR*-/- embryos, suggesting that their atypical upregulation very early in development could affect proper embryo differentiation and contribute to the deficient ability of *AhR*-/- mice to sustain implantation and in-utero survival (Abbott et al., 1999; Fernandez-Salguero et al., 1995; Peters and Wiley, 1995). The pro-differentiation role of AhR in embryogenesis is also supported by its own regulation during the process. AhR levels increased with differentiation and, interestingly, its location was mainly restricted to the blastomeres that differentiate to form the trophoectoderm, indicating that AhR may exert a differential regulatory function limiting pluripotency in those blastomeres that will generate extraembryonic tissues. In this sense, a central regulator such as OCT4, showed an opposite expression pattern to that of AhR, again supporting its repressive role in pluripotency. The crosstalk between OCT4 and AhR has been also suggested from studies using stem-like cancer cells which proposed a reciprocal suppression between AhR and such pluripotency factor (Cheng et al., 2015; Song et al., 2002).

In the morulae, it is known that the first asymmetric division is determinant for embryonic differentiation, and that in the formed blastocyst a second differentiating wave gives rise to two types of cells in the ICM. The fact that AhR expression was modulated during these processes, together with previous studies that link embryo differentiation to the Hippo pathway, lead us to think that AhR could act through Hippo in the phenotype observed. Our preliminary data indicate that nuclear YAP levels can be modulated by AhR producing a more differentiated status in NTERA-2 cells (Morales-Hernández et al., unpublished results). In this work, we have shown that TE cell fate seems to be activated by nuclear YAP in an AhR-dependent manner, and thus, AhR and YAP co-localized in the nucleus of external blastomeres eventually differentiating to the trophectoderm lineage. The fact that AhR-null blastocysts had OCT4 expression but lacked nuclear YAP in external blastomeres, suggest that AhR deficiency may result in a failure to link polarity to transcription factors that lead to differentiation through Hippo signalling.

One characteristic of pluripotent cells is their low levels of oxidative phosphorylation and their preferred glycolytic ATP synthesis. Up-regulation of glycolysis preceedes the reactivation of pluripotent markers (Shyh-Chang et al., 2013). The differences that we have observed in differentiation markers through embryo development were correlated with their metabolic status. Lack of AhR reduced mitochondrial activity and maintained a predominant glycolytic metabolism. During differentiation, metabolic pathways are modulated according to the needs of the embryo. Our results are in agreement with those hypothesis since *AhR*-/- embryos overexpressed threonine dehydrogenase (TDH), which is an enzyme responsible for providing metabolites generated from Thr that are specifically used for stem cell self-renewal. Therefore, lack of AhR probably causes a metabolic state in the embryos that corresponds to a lower differentiation state. In summary, AhR has relevant functions in embryonic development adjusting the expression of signaling pathways that control pluripotency and diferentiation. Under low AhR levels, a defective differentiation status may compromise completion of the embryo developmental program, implantation and survival.

## MATERIALS AND METHODS

### Embryo collection

C57BL/6N wild-type (*AhR+/+*) and AhR-null (*AhR*-/-) mice were kept under 12 h light/dark cycle and had free access to food and water. 4 to 7 weeks old females were injected with 7.5 IU Pregnant Mare’s Serum followed 48 h later by 5 IU i.p. injection of human chorionic gonadotropin (hCG). Females were sacrificed at the indicated developmental stages and the oviducts/hemiuterus were collected in PBS and flashed for embryo collection. Embryos were isolated using a stripper (Origio). All work involving mice has been performed in accordance with the National and European legislation (Spanish Royal Decree RD53/2013 and EU Directive 86/609/CEE as modified by 2003/65/CE, respectively) for the protection of animals used for research. Experimental protocols using mice were approved by the Bioethics Committee for Animal Experimentation of the University of Extremadura (Registry 109/2014) and by the Junta de Extremadura (EXP-20160506-1). Mice had free access to water and rodent chow.

### Gene expression analysis

Total RNA was isolated from mouse embryos using the pico pure RNA isolation Kit (Thermo Fisher) and purified following the manufacturer’s instructions. Reverse transcription was performed using random priming and the iScript Reverse Transcription Super Mix (Bio-Rad). Real-time PCR was used to quantify the mRNA expression of *AhR, Nanog, Oct4, Sox2, Lats1, Lats2, β-catenin, Scl2a1, Scl2a2, Hexokinase, Cdx2, Gata3, TDH*. Reactions were done using Luna Master Mix (New England Biolabs) in a step one thermal cycler (Applied Biosystems) essentially as described (Rey-Barroso et al., 2013). The expression of *β*-*Actin* was used to normalize gene expression (ΔCt) and 2^−ΔΔCt^ was applied to calculate changes in RNA levels with respect to control conditions. Primer sequences used are indicated in supplementary Table S1.

### Whole mount immunofluorescence

Each embryo group was independently fixed in 3.5% paraformaldehyde for 15 min at room temperature. The zona pellucida was removed by incubation in Tyrode’s acid solution for 15-20 s at 37°C. Embryos were blocked in PBS containing 1% BSA and 0.1 M glycine for 2.5 h followed incubation in blocking solution with antibodies against NANOG, OCT4, AhR, YAP, pYAP overnight at 4°C. Following washings, an Alexa-633, 488 or 550 labeled secondary antibodies was added for 2 h at 4°C. Samples were further washed and incubated with Hoechst to stain cell nuclei. Embryos were transferred to Ibidi chambers and analyzed using an Olympus FV1000 confocal microscope.

### Magnetic-activated Cell Sorting

Inner cell mass (ICM) and trophectoderm (TE) blastomeres were separated using concanavalin and MACS microbeads essentially as described (Ozawa and Hansen, 2011). Blastocysts at 3,5 d.p.c were harvested and incubated in acidic Tyrode’s solution to remove the zona pellucida. Samples were washed three times in MACS buffer [DPBS with 0.5% (w/v) BSA and 2 mM ethylenediaminetetraacetic acid (EDTA), pH 7.2] and incubated for 10 min with concanavalin A conjugated-FITC (Sigma-Aldrich, ConA-FITC, 1 mg/ml in MACS buffer). Following three washes in MACS buffer, blastocysts were incubated in PBS containing 1 mM EDTA for 5 min followed by incubation in 0.05% (w/v) trypsin-0.53 mM EDTA solution (Invitrogen) for 10 min at 37°C. Groups of 15-20 blastocysts were disaggregated into single blastomeres by pipetting with an stripper (Origen) under a dissecting microscope. Blastomeres were transferred into PBS containing 1 mM EDTA and 10% (v/v) fetal bovine serum to stop the reaction. Samples were then washed in MACS buffer by centrifugation at 500 x g for 5 min and resuspended in 110 μl of MACS buffer. Disaggregated blastomeres in solution were incubated with 10 μl of magnetic microbeads conjugated to mouse anti-FITC (Miltenyi Biotec) for 15 min on ice. Following two washes by centrifugation at 500 x g for 5 min, samples were resuspended in 500 μl MACS buffer and passed through MACS separation columns (Miltenyi Biotec) attached to a magnetic board (Spherotech). The FITC-negative fraction (ICM) was eluted by three 500 μl MACS buffer washes followed by FITC positive (TE) elution by removing the MACS separation column from the magnetic board and washing three times with 500 μl MACS buffer.

### TMRM and Mitotracker staining

The embryos were arranged in staining solution made with 10 μL of 100 μM stock solution Tetramethylrhodamine (Invitrogen) in 10 mL of KSOM medium (EmbrioMax, Millipore) or 100 nM of Mitotracker green (Cell Signalling). Embryos were placed in IBIDI plates in a 5% CO_2_ incubator at 37°C for 30 min for TMRM staining and 20 min for mitotracker. Then, embryos were washed in PBS twice and analyzed by confocal microscopy.

### Mitochondrial potential measurement using the JC10 Kit

Pools of 25 embryos were placed in a 96 plate well, and processed following the non-adherent cell protocol recommended by the manufacturer. An aliquot of 50 μl of JC-10 dye loading solution was added per well and the embryos were incubated in a 5% CO_2_ incubator at 37 °C for 30 min. Then, 50 μl of assay buffer were added and fluorescence intensity was monitored in a fluorescence multiwell plate reader using excitation wavelength 490 nm and emission wavelength 525 nm. For ratio analysis, signals were also recorded at excitation wavelength 540 nm and emission wavelength 590 nm. The red/green fluorescence intensity ratio was used to determine the mitochondrial membrane potential (MMP).

### Hexokinase activity measurement

Groups of 20 blastocysts were disaggregated by incubation in 0.05% (w/v) trypsin solution containing 0.53 mM EDTA for 10 min at 37°C. Samples were centrifuged, washed twice in PBS containing 1 mM EDTA and 10% (v/v) fetal bovine serum and once in PBS. Single blastomeres were resuspended in hexokinase (HK) assay buffer and homogenized through passage by a 30 g syringe. Homogenized samples were used in the pico-probe Hexokinase activity assay kit (Biovision) following manufacturer’s indications.

### Statistical analyses

GraphPad Prism 6.0 software (GraphPad) was used to perform comparison between experimental conditions. Student’s t-test (unpaired two-sided) was used to analyze differences between two experimental groups. For three or more experimental conditions data was analyzed using ANOVA. Data are shown as mean ± SD. Differences were considered significant at p*<0.05; p**<0.01; p***<0.001. Data analyses are indicated in the Figure Legends.

## ACKNOWLEDGMENTS

The Servicio de Técnicas Aplicadas a las Biociencias (STAB) of the Universidad de Extremadura are greatly acknowledged for their technical support.

## REFERENCES

Abbott, B.D., Schmid, J.E., Pitt, J.A., Buckalew, A.R., Wood, C.R., Held, G.A., and Diliberto, J.J. (1999). Adverse reproductive outcomes in the transgenic Ah receptor-deficient mouse. Toxicol. Appl. Pharmacol. 155, 62–70.

Artus, J., Piliszek, A., and Hadjantonakis, A.K. (2011). The primitive endoderm lineage of the mouse blastocyst: sequential transcription factor activation and regulation of differentiation by Sox17. Dev. Biol. 350, 393–404.

Bedzhov, I., Graham, S.J., Leung, C.Y., and Zernicka-Goetz, M. (2014). Developmental plasticity, cell fate specification and morphogenesis in the early mouse embryo. Philosophical Transactions of the Royal Society B: Biological Sciences 369, 20130538.

oyer, L.A., Lee, T.I., Cole, M.F., Johnstone, S.E., Levine, S.S., Zucker, J.P., Guenther, M.G., Kumar, R.M., Murray, H.L., Jenner, R.G., et al. (2005). Core transcriptional regulatory circuitry in human embryonic stem cells. Cell 122, 947–956.

Boyer, L.A., Mathur, D., and Jaenisch, R. (2006). Molecular control of pluripotency. Curr. Opin. Genet. Dev. 16, 455–462.

Chazaud, C., and Yamanaka, Y. (2016). Lineage specification in the mouse preimplantation embryo. Development 143, 1063–1074.

Cheng, J., Li, W., Kang, B., Zhou, Y., Song, J., Dan, S., Yang, Y., Zhang, X., Li, J., and Yin, S. (2015). Tryptophan derivatives regulate the transcription of Oct4 in stem-like cancer cells. Nature communications 6, 7209.

Fernandez-Salguero, P., Pineau, T., Hilbert, D.M., McPhail, T., Lee, S.S., Kimura, S., Nebert, D.W., Rudikoff, S., Ward, J.M., and Gonzalez, F.J. (1995). Immune system impairment and hepatic fibrosis in mice lacking the dioxin-binding Ah receptor. Science 268, 722–726.

Fleming, T.P. (1987). A quantitative analysis of cell allocation to trophectoderm and inner cell mass in the mouse blastocyst. Developmental biology 119, 520–531.

Frum, T., and Ralston, A. (2015). Cell signaling and transcription factors regulating cell fate during formation of the mouse blastocyst. Trends Genet. 31, 402–410.

Johnson, M.H., and Ziomek, C.A. (1981). The foundation of two distinct cell lineages within the mouse morula. Cell 24, 71–80.

Ko, C.I., and Puga, A. (2017). Does the Aryl Hydrocarbon Receptor Regulate Pluripotency? Curr Opin Toxicol 2, 1–7.

Laiosa, M.D., Tate, E.R., Ahrenhoerster, L.S., Chen, Y., and Wang, D. (2015). Effects of Developmental Activation of the Aryl Hydrocarbon Receptor by 2,3,7,8-Tetrachlorodibenzo--dioxin on Long-Term Self-Renewal of Murine Hematopoietic Stem Cells. Environ. Health Perspect.

Leung, C.Y., and Zernicka-Goetz, M. (2013). Angiomotin prevents pluripotent lineage differentiation in mouse embryos via Hippo pathway-dependent and-independent mechanisms. Nature communications 4, 2251.

Manzanares, M., and Rodriguez, T.A. (2013). Development: Hippo signalling turns the embryo inside out. Current Biology 23, R559–R561.

Morales-Hernandez, A., Gonzalez-Rico, F.J., Roman, A.C., Rico-Leo, E., Alvarez-Barrientos, A., Sanchez, L., Macia, A., Heras, S.R., Garcia-Perez, J.L., Merino, J.M., et al. (2016). Alu retrotransposons promote differentiation of human carcinoma cells through the aryl hydrocarbon receptor. Nucleic Acids Res 44, 4665–4683.

Morales-Hernandez, A., Nacarino-Palma, A., Moreno-Marin, N., Barrasa, E., Paniagua-Quinones, B., Catalina-Fernandez, I., Alvarez-Barrientos, A., Bustelo, X.R., Merino, J.M., and Fernandez-Salguero, P.M. (2017). Lung regeneration after toxic injury is improved in absence of dioxin receptor. Stem Cell Res 25, 61–71.

Moreno-Marin, N., Barrasa, E., Morales-Hernandez, A., Paniagua, B., Blanco-Fernandez, G., Merino, J.M., and Fernandez-Salguero, P.M. (2017). Dioxin Receptor Adjusts Liver Regeneration After Acute Toxic Injury and Protects Against Liver Carcinogenesis. Sci Rep 7, 10420.

Morris, S.A., Teo, R.T., Li, H., Robson, P., Glover, D.M., and Zernicka-Goetz, M. (2010). Origin and formation of the first two distinct cell types of the inner cell mass in the mouse embryo. Proceedings of the National Academy of Sciences 107, 6364–6369.

Mulero-Navarro, S., and Fernandez-Salguero, P.M. (2016). New Trends in Aryl Hydrocarbon Receptor Biology. Front Cell Dev Biol 4, 45.

Ozawa, M., and Hansen, P.J. (2011). A novel method for purification of inner cell mass and trophectoderm cells from blastocysts using magnetic activated cell sorting. Fertility and sterility 95, 799–802.

Paramasivam, M., Sarkeshik, A., Yates III, J.R., Fernandes, M.J., and McCollum, D. (2011). Angiomotin family proteins are novel activators of the LATS2 kinase tumor suppressor. Molecular biology of the cell 22, 3725–3733.

Peters, J.M., and Wiley, L.M. (1995). Evidence that murine preimplantation embryos express aryl hydrocarbon receptor. Toxicol. Appl. Pharmacol. 134, 214–221.

Rey-Barroso, J., Colo, G.P., Alvarez-Barrientos, A., Redondo-Munoz, J., Carvajal-Gonzalez, J.M., Mulero-Navarro, S., Garcia-Pardo, A., Teixido, J., and Fernandez-Salguero, P.M. (2013). The dioxin receptor controls beta1 integrin activation in fibroblasts through a Cbp-Csk-Src pathway. Cell Signal 25, 848–859.

Rico-Leo, E.M., Moreno-Marin, N., Gonzalez-Rico, F.J., Barrasa, E., Ortega-Ferrusola, C., Martin-Munoz, P., Sanchez-Guardado, L.O., Llano, E., Alvarez-Barrientos, A., Infante-Campos, A., et al. (2016). piRNA-associated proteins and retrotransposons are differentially expressed in murine testis and ovary of aryl hydrocarbon receptor deficient mice. Open Biol 6.

Roman, A.C., Carvajal-Gonzalez, J.M., Merino, J.M., Mulero-Navarro, S., and Fernandez-Salguero, P.M. (2018). The aryl hydrocarbon receptor in the crossroad of signalling networks with therapeutic value. Pharmacol. Ther. 185, 50–63.

Shyh-Chang, N., Daley, G.Q., and Cantley, L.C. (2013). Stem cell metabolism in tissue development and aging. Development 140, 2535–2547.

Song, J., Clagett-Dame, M., Peterson, R.E., Hahn, M.E., Westler, W.M., Sicinski, R.R., and DeLuca, H.F. (2002). A ligand for the aryl hydrocarbon receptor isolated from lung. Proceedings of the National Academy of Sciences 99, 14694–14699.

Wang, J., Alexander, P., Wu, L., Hammer, R., Cleaver, O., and McKnight, S.L. (2009). Dependence of mouse embryonic stem cells on threonine catabolism. Science 325, 435–439.

Wang, Q., Kurita, H., Carreira, V., Ko, C.I., Fan, Y., Zhang, X., Biesiada, J., Medvedovic, M., and Puga, A. (2016). Ah Receptor Activation by Dioxin Disrupts Activin, BMP, and WNT Signals During the Early Differentiation of Mouse Embryonic Stem Cells and Inhibits Cardiomyocyte Functions. Toxicol Sci 149, 346–357.

